# A function of Spalt major as a sequence-specific DNA binding transcription factor mediates repression of *knirps* in the *Drosophila* wing imaginal disc

**DOI:** 10.1101/2023.09.04.556233

**Authors:** Cristina M. Ostalé, Alicia del Prado, Nuria Esteban, Ana López-Varea, Jose F. de Celis

**Affiliations:** Centro de Biología Molecular “Severo Ochoa” CSIC and Universidad Autónoma de Madrid Madrid 28049 Spain

**Keywords:** Spalt proteins, *knirps* regulation, gene expression, vein patterning

## Abstract

The Spalt transcriptional regulators participate in a variety of cell fate decisions during multicellular development. Vertebrate Spalt proteins have been mostly associated to the organization of heterochromatic regions, but they also contribute regulatory functions through binding to A/T rich motives present in their target genes. The developmental processes in which the *Drosophila spalt* genes participate are well known through genetic analysis, but the mechanism by which the Spalt proteins regulate transcription are still unknown. Furthermore, despite the prominent changes in gene expression associated to mutations in the *spalt* genes, the specific DNA sequences they bind are also unknow. In this contribution we describe the analysis of a DNA fragment present in the regulatory region of the *knirps* gene. Spalt proteins are candidate repressors of *knirps* expression during the formation of the venation pattern in the wing disc, and here we identify a minimal conserved 30bp sequence that binds to Spalt major both *in vivo* and *in vitro*. This sequence mediates transcriptional repression in the central region of the wing blade, constituting the first confirmed case of a direct regulatory interaction between Spalt major and its target DNA in *Drosophila*.

## INTRODUCTION

The Spalt proteins (Sal) are transcriptional regulators that play key roles in a variety of cell fate decisions during multicellular development (de Celis and Barrio, 2009). Their structure is very much conserved from *C. elegans* to humans, and their main characteristic motives are pairs of C2H2 Zn finger domains distributed along the length of the protein (de Celis and Barrio, 2009; Alvarez et al., 2021). Human *sal* genes, named *sal-like* genes (SALL) have been the subject of considerable research, because mutations in two of then are causative of severe congenital disorders, the Towles-Brook disease (SALL1; Powell and Michaelis, 1999; Kohlhase, 2000) and the Okihiro syndrome (SALL4; Borozdin et al., 2004). In addition, SALL genes are increasingly found as reliable markers of oncogenic malignancy, because their inappropriate expression in many adult tissues promotes a stem cell-like phenotype that favors tumoral growth and metastasis (Alvarez et al., 2021). The pathologies associated to mutations or inappropriate expression of human SALL genes are most likely related to the functions that these genes play during normal development, which have been mostly characterize in murine and *Drosophila* models (Sweetman and Münsterberg, 2006; de Celis and Barrio, 2009). A common aspect of the functional requirements of *sal* genes is their participation during embryonic development in multitude of developmental processes occurring during organogenesis and stem cell formation. In many of these processes *sal* function is required to promote alternative cell fates acting as transcriptional regulators.

The *sal* genes were first identified in *Drosophila*, where the two *sal* genes present, *spalt major* (*salm*) and *spalt related* (*salr*) form part of a gene complex (Barrio et al., 1996; de Celis et al., 1996). In this organism, the function of these genes is required for neural development (Cantera et al., 2002), the assignation of cell identities in the mesoderm and peripheral neural system (Rusten et al., 2001; Schönbauer et al., 2011), the formation of the dorsal tracheal truck and the posterior spiracles (Kühnlein and Schuh; 1996; Hu and Castelli-Gair, 1999) and the formation of the wing, among others (Grieder et al., 2009; Organista and de Celis, 2012). In some of these processes it has been shown that Sal proteins and genes are part of diverse gene regulatory networks, where their expression is regulated by different transcription factors acting on shared enhancers affecting *salm* and *salr* expression (Barrio et al., 1999; Martin et al., 2014). In turn, the Salm and Salr proteins regulate other downstream genes, forming transcriptional cascades resulting in cell differentiation or territorial specification. It has proven very difficult, however, to determine direct binding of Sal proteins to the DNA of candidate downstream genes, as there is no sequence binding motive describe for these proteins. In this manner, the only motive identified as bound by a Sal protein in *Drosophila* is a TATGAAAT sequence present in the *s15* chorion gene (Shea et al. 1990) that is bound by a fragment of the Salr protein (Barrio et al., 1996). More recently, it has been shown that mouse SALL1, SALL3 and SALL4 bind preferentially AT rich regions containing AATA tetranucleotides (Ru et al., 2022), and that human SALL4 preferentially uses AA[A/T]TAT[T/G][A/G][T/A] motives as binding sequences to regulate its target genes in cancer cells (Kong et al., 2021).

The *Drosophila sal* genes play several roles during the growth and patterning of the wing imaginal disc. This epithelial tissue grows during larval development and became progressively patterned by the restricted expression of several transcription factors, which domains of expression label developing territories first, and specific pattern elements, such as the precursors of the sensory organs and veins, later. The function of *salm/salr* in the wing disc is first required to specify the more proximal territory of the disc, which will give rise to the thorax (Grieder et al., 2009). Later in this territory, *salm/salr* are required for the patterning of several sensory organs (de Celis et al., 1999). In the developing wing blade, the *salm/salr* genes are expressed under regulation by the BMP pathway in a central territory of the presumptive wing blade (de Celis et al., 1996), where they are required to promote cell division and survival and to specify the position of the two peripheral longitudinal veins, the L2 in the anterior compartment and the L5 in the posterior compartment (Organista and de Celis, 2012). The specification of the L2 vein involves regulatory interactions between several transcription factors encoded by the genes *optix*, *salm/salr* and *knirps* (*kni*) (Lunde et al., 1998; de Celis and Barrio, 2000). Thus, the presumptive L2 territory corresponds to the domain of *kni* expression, and its anterior and posterior limits are determined by repression mediated by Optix and Salm/Salr, respectively (Martin et al., 2017).

Here we show that the regulation of *kni* involves direct repression mediated by the Salm/Salr proteins. This repression occurs through a sequence of 59 bp located 11.800 bp from the *kni* transcription start site. This sequence is highly conserved in *Drosophilids*, and we show that Salm binds to this sequence *in vivo* (ChIP) and *in vitro* (EMSA). To our knowledge, this sequence, which confers Salm/Salr repression to heterologous activators, is the first confirmed Sal direct target identified in *Drosophila*. These results define a precise mechanism of Salm repression on *kni* that links the BMP gradient to the positioning of the anterior limit of expression of a gene involved in the development of the longitudinal L2 vein.

## MATERIALS AND METHODS

### Drosophila strains

We use the Gal4 drivers *sal^EPv^-Gal4* (Cruz *et al*. 2009), *hh-Gal4* and *nub-Gal4* (Calleja *et al*. 1996), the UAS-RNAi lines *UAS-salm-i* (VDRC3029 and BL60462), *UAS-salr-i* (VDRC28386 and BL29549), from the Vienna *Drosophila* Resource Center (VDRC) and Bloomington *Drosophila* Stock Center (BDSC). An endogenous tagged Salm version, *Salm-HA-3xFlag* (F1049 line) was a gift from Frank Schnorrer. Flies were raised at 25°C (unless otherwise stated) and 55% relative humidity in fly medium containing Glucose (50gr/L), Agar (7.86 gr/L), wheat flour (35,7 gr/L), yeast (71,4 gr/L), Methylparaben (2,8ml/L) and Propionic acid (4,3 ml/L).

#### Generation of GFP reporter constructs

All transformants described below were generated by ϕC31 site-specific integration in the 3R chromosomic arm in a y[1] M{RFP[3xP3.PB] GFP[E.3xP3]=vas-int.Dm}ZH-2A w[*]; M{3xP3-RFP.attP}ZH-86Fb background (Bischof et al., 2007).

##### kni regulatory regions

We cloned a 691 bp fragment from the *kni* regulatory region (kni- RA) in pHP-Dest-eGFP (Plásmido #24566 Addgene) (Boy et al., 2010) using LR-Clonasa IITM (Invitrogen). This fragment was first amplified by PCR from kniRR (Martin et al., 2017) using the oligonucleotides *kniRR* Fw and *kniRA* Rv (Supple. Table 1) and cloned in p-ENTR/D-TOPO (Invitrogen). Cloning was confirmed by PCR using the oligonucleotides M13 Fw and kniRA Rv (Supple. Table 1). The fragment *kni-RA*Δ*4* was amplified from *kni- RA* using the oligonucleotides *kni-RA*Δ*4* Fw and *kni-RA* Rv (Supple. Table 1). The fragments *kni-RA*Δ*5* was amplified from *kni-RA*Δ2 (Martin et al., 2019) using the oligonucleotides *kni-RA*Δ*2* Fw and *kni-RA*Δ*5* Rv (Supple. Table 1). The fragments *kni- RA*Δ*4* and *kni-RA*Δ*5* were also cloned in p-ENTR/D-TOPO and then transferred to pHP- Dest-eGFP using LR-Clonasa IITM.

##### Ser constructs

The constructs *wingSer-kniRA-*Δ3, *wingSer-kniRA-*Δ4 and *wingSer-60sal* were made from the fragments *kniRA-*Δ*3* (Martin et al., 2017)*, kniRA-*Δ*4* and *60sal* cloned in pENTR/D-TOPO containing the wing regulatory region of *Serrate* (wing-Ser; Yan et al., 2004) by recombinant PCR (Supple. Fig. 1). The sequence of the different oligonucleotides using in these PCRs are described in Suppl. Table 1.

**Figure 1.**
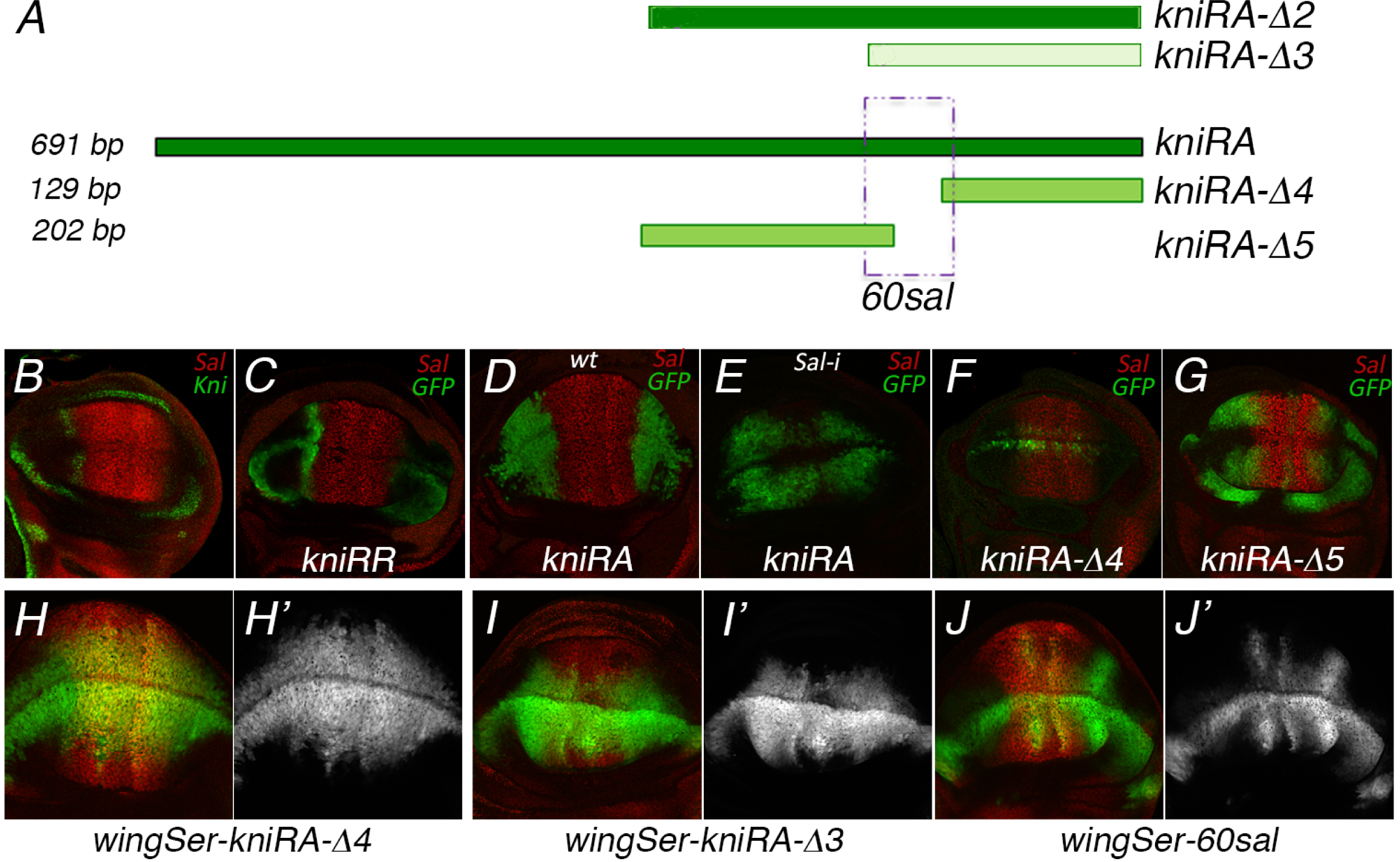
Identification of *60sal*, a sequence that confer repressor activity to the *kni* L2 vein enhancer (A) Schematic representation of the 691 bp region of kni (*kniRA*) and different fragments of this sequence analyzed in Martin et al., 2017 (*kniRA-*Δ*2* and *kniRA-*Δ*3*) and in this manuscript (*kniRA-*Δ*4*, *kniRA-*Δ*5* and *60sal*). Green color indicates expression of GFP when its coding region is fused to these regulatory fragments. (B) Expression of Kni (green) and Salm (red) in a mature third instar wing disc. (C-D) Expression of Salm (red) and GFP in third instar wing discs from *kni-RR-GFP* and *kni-RA-GFP* larvae. (E) Expression of Salm (red) and GFP in third instar wing discs from *sal^EPv^-Gal4/UAS-salm-RNAi; UAS- salr-RNAi/ kni-RA-GFP* larvae. (F-G) Expression of Salm (red) and GFP in third instar wing discs of *kni-RA-* Δ*4-GFP* and *kni-RA-* Δ*5-GFP* genotypes. (H-J) Expression of Salm (red) and GFP in third instar wing discs from larvae of *wingSer-kni-RA-*Δ*4-GFP* (H) *wingSer-kni- RA-*Δ*3-GFP* (I) and *wingSer-60sal-GFP* (J) genotypes. Individual green channels are shown in white in H’-J’.

##### UAS-60sal constructs

the sequence *60sal* was cloned in pC4PLZ (a gift from Gerardo Jiménez) by recombinant PCR using the oligonucleotides UAS Fw, UAS-(60sal) Rv, (UAS)- *60sal* Fw and *60sal* Rv (Supp. Fig. 1). The constructs *UAS-60sal-*Δ*5, UAS-60sal-*Δ*3-1* y *UAS-60sal-*Δ*3-2* were generated from pENTR/D-TOPO containing UAS(5x)-60sal(1x) with a KpnI site inserted at the 5’ end of Δ*3-1* or Δ*3-2* (Quick Change Site-Directed Mutagenesis from Stratagene) using the oligonucleotides Δ*3-1-KpnI* Fw, Δ*3-1-KpnI* Rv, Δ*3-2-KpnI* Fw and Δ*3-2-KpnI* Rv (Supple. Table 1). The constructs 60sal-MUT1, 60sal- MUT2 and 60sal-MUT3 were generated from pENTR/D-TOPO-UAS(5x)-60sal(1x) using Pfu polymerase and Quick Change Site-Directed Mutagenesis (Stratagene). Sequences of 6 nucleotides were changed C to A and G to T using the oligonucleotide pairs 60sal- MUT1 Fw and 60sal-MUT1 Rv, 60sal-MUT2 Fw and 60sal-MUT2 Rv, 60sal-MUT3 Fw and 60sal-MUT3 Rv. The constructs *UAS(5x)-60salMUT(1x)* were used to generate the corresponding *UAS(5x)-60salMUT(2x)* by recombinant PCR using a workflow similar to that show in Suppl. Fig. 1B and the oligonucleotides *UAS* Fw, 60sal db Rv, *60salMUT* Fw and *60salMUT* Rv (Supple. Table 1).

### Salm CDS cloning, protein expression and EMSA assays

#### pDest17-SalmCDS

The Salm cDNA was amplified from pBlueScript II SK containing the *salm* cDNA (a gift from Rosa Barrio) using the oligonucleotides *salCDS Fw* and *salCDS Rv* and cloned in pENTR/D-TOPO (Invitrogen). The *salm.CDS* cassette was transferred to pDest17 by LR-Clonasa IITM (invitrogen), and the positive clones were confirmed by DNA sequencing using the oligonucleotide salCDC check (Supple. Table 1). The construct pDest-17-SalmCDC allows the expression in *E. coli* BL21 of the Salm protein fused to a 6 Histidine tail using the T7 promoter. The Salm protein was purified from protein extracts of *E. coli* BL21 transformed with pDest17-SalmCDS and pLYSs. Protein expression was induced by IPTG (0,6mM 24h room temperature) after an overnight culture grown in SOC (LB, glucosa 2mM) with Ampiciline (80µg/ml) and Chloramphenicol (80µg/ml). For Salm protein purification from bacterial lysates (lisozima 0,5mg/mL, Imidazol 10mM and protease inhibitors *cOmplete™ Protease Inhibitor Cocktail tablets* from Roche) we used 3 rounds of 3 minutes sonication with 15 s on/off cycles. The samples were then treated with DNAse (5μg/mL) and RNAse (10μg/mL) for 15 min on ice, and after standard treatments the Salm protein was eluted from Histidine columns and visualized by western blot using mouse anti-His (1:1000; Dianova) and anti-Salm (1:200; Barrio et al., 1999).

The interaction of Salm proteins with the primer–template structure was assayed using as substrate the 5’-labeled oligonucleotides *60sal S* and *60sal AS* for the entire *60sal* fragment, and the pairs *60salC S*/*60salC AS* and *60salNC S*/*60salNC A*S (Supple. Table 1). For a negative control we used the independent oligonucleotides *sp1 S* and *sp1 AS* (Supple. Table 1). Primer oligonucleotides were 5’-labeled with ^32^P using [g-^32^P]ATP (10 mCi) and T4PNK. The hybrid molecule was obtained by annealing the primer to template (1:2 ratio) in the presence of 50 mM Tris- HCl pH 7.5 and 0.2 M NaCl, with heating to 90 °C for 10 min before slowly cooling to room temperature overnight. The [g-^32^P]ATP (3,000 Ci/mmol) was supplied by PerkinElmer. T4 polynucleotide kinase (T4PNK) was purchased from New England Biolabs. The incubation mixture contained, in a final volume of 20 µl, 12 mM Tris-HCl, pH 7.5, 20 mM ammonium sulphate, 0.1 mg/ml BSA, 0,16 ng of the indicated template/primer DNA molecule and the purified Salm protein (0, 100 ng, 1 µg or 10 µg). After incubation for 5 min at 4 °C, the samples were subjected to electrophoresis in precooled 4% (w/v) polyacrylamide gels (acrylamide/bis-acrylamide 80:1, w/w) containing 12 mM Tris-acetate, pH 7.5 and run at 4 °C in the same buffer at 8 V/cm (Carthew *et al.,* 1985). After autoradiography, Salm/DNA stable interaction was detected as a shift (retardation) in the migrating position of the labelled DNA.

### Immunocytochemistry

Imaginal discs were dissected in PBS (NaCl 138mM; KCl 3mM; Na2HPO4 8,1 mM; KH2PO4 1.5 mM) and fixed 20 min at RT in PBT (PBS/Tritón X-100 0,3%) with 4% formaldehyde. The discs were pre-incubated at least for 30 min in PBT-BSA (1% BSA from Sigma), and incubated with primary antibodies ON at 4 C. We used the following primary antibodies: Rat and Rabbit anti-Salm (1:200; Barrio et al., 1999) and Guinea pig anti-Kni 1:500; Kosman et al., 1998). Primary antibodies were rinsed with PBT (3x 15 mins) and the discs incubated with alexa-conjugated secondary Rat, Rabbit, Mouse or Guineapig secondary antibodies (ThermoFisher). In most staining’s we used DAPI (1:4000; Merk) to reveal DNA. After several rinses in PBT the wing discs were mounted in Vectashield (Vector Laboratories), and visualized in Zeiss confocal microscopes LSM510, LSM710 or LSM900. All images were processed with ImageJ2 v2.3.0/1.53q (ImageJ) and Photoshop v24.4.1 (Adobe^TM^).

### Salm chromatin immunoprecipitation

We made chromatin immunoprecipitation followed by sequencing (ChIP-Seq) from wing imaginal discs of Salm-3xFlag genotype (strain F1049, courtesy of Dr. Frank Schnorrer) using anti-Flag antibodies (Sigma-Aldrich). The samples from anti-Flag immunoprecipitated chromatin and controls were sequenced at the Centre for Genomic Regulation, Barcelona, Spain). The ChIP-Seq experiments will be described in full elsewhere (CMO and JFdC, in preparation).

## RESULTS and DISCUSSION

### Regulation of *kni* expression in the wing disc

The L2 provein is defined as the territory of *kni* expression in the anterior compartment of the *Drosophila* wing blade (Fig. 1B). The regulation of *kni* expression in this domain involves a general activation mechanism and the function of two repressors, Optix and Salm/Salr, which set the anterior (Optix) and the posterior (Salm/Salr) limits to the *kni* expression domain (Fig. 1A-C; Lunde et al., 2003; Martin et al., 2017; Xu et al., 2017). We started the analysis of the *kni* regulatory region with a 691 bp fragment (*kni-RA*) identified by Lunde et al. (2003). This fragment, originally defined as *kni* “Activation domain” is not able to drive reporter expression in the territory where Salm and Salr are expressed (Fig. 1A, D), indicating that contains the sites mediating *kni* repression. Loss of Salm/Salr function in this territory results in reporter expression in the central domain of the wing disc (Fig. 1E), confirming that these proteins mediate repression in these cells. The expression of the reporter is not affected by proximal deletions of up to 330 bp from 5’ terminus (*kniRA-*Δ*1* and *kniRA-*Δ*2*; Martin et al., 2017). However, deletion of the adjacent 178 bp (*kniRA-*Δ*3*; Fig. 1A), entirely removes reporter expression (Martin et al., 2017), indicating that this sequence includes a region required for *kni* activation. We generated two additional reporter constructs of 129 (*kniRA-*Δ*4*; Fig. 1F) and 202 (*kniRA-*Δ*5*; Fig. 1G) bp in length, aiming to preserve the sequences required for activation. We found that *kniRA-*Δ*5* retains both the activating sequences and the DNA mediating repression in the central domain of the wing (Fig. 1G). In contrast, *kniRA-*Δ*4* retains some activation of the reporter in regions close to the dorso-ventral boundary, but losses reporter repression in the central domain of the wing (Fig. 1F).

The close proximity of activating and repressing sequences included in these constructs prevents the analysis of repression, and consequently we shifted to fusion constructs in which the activating sequences are provided by heterologous DNA. We first made fusion constructs containing as activating sequences an enhancer of the Serrate gene (*wingSer*; Yan et al., 2004). The fusion of *wingSer* to *kniRA*Δ*4*, which lack the repression sequences of *kni*, causes reporter expression in the entire wing blade (Fig. 1H-H’). In contrast, the fusion of *wingSer* to the longer fragment *kniRA*Δ*3* results in a consistent loss of reporter expression in the central region of the wing blade (Fig. 1I-I’). When we fused 59 bp including the region of *kniRA*Δ*3* that is missing in *kniRA*Δ*4* (*60sal* fragment; Fig. 1A) to *wingSer*, we found an even more consistent repression of reporter expression in the central domain of the wing blade (Fig. 1J-J’), suggesting that this small DNA sequence confers some repression to the *wingSer*-*GFP* construct. We decided to pursue the analysis of this 59 bp fragment (*60sal*) by generating novel reporter constructs containing an array of UAS activating sequences fused to one or two copies of the *60sal* sequence. We first analyzed the effect of the *60sal* silencer in the entire wing blade by driving the expression of GFP from the UAS sequences with *nubbin-Gal4* (*nub- Gal4*). Compared to the control (*5xUAS*; Fig. 2A), the introduction of one copy of *60sal* (*5xUAS/1x60sal*) causes a weak reduction in reporter expression in the central domain of the wing, where the Salm/Salr proteins are expressed (Fig. 2B). An additional copy of *60sal* (*5xUAS/2x60sal*) entirely prevents reporter expression in the wing region where Salm/Salr are present (Fig. 2C). Finally, when only one copy of activating sequences and repressive sequences are present (*1xUAS/1x60sal*), we also observe a moderate repression in the central region of the wing (Fig. 2D). We obtained similar results using other Gal4 drivers, such as *sal-Gal4* (Fig. 2E-F) and *hh-Gal4* (Fig. 2G-H), confirming that the *60sal* fragment contains regulatory elements mediating the repression of the reporter constructs in any territory of the wing blade where Salm/Salr are expressed. Furthermore, our results indicate that the final outcome of reporter expression depends on the ratio of activating sequences (*UAS* copies) and repressive ones (*60sal*). Interestingly, the repressive action of Salm/Salr appears restricted to the wing blade, because our reporters are insensitive to the presence of these proteins in other tissues such as the trachea (Fig. 2I-J) and the ring gland (Fig. 2K-K’).

**Figure 2.**
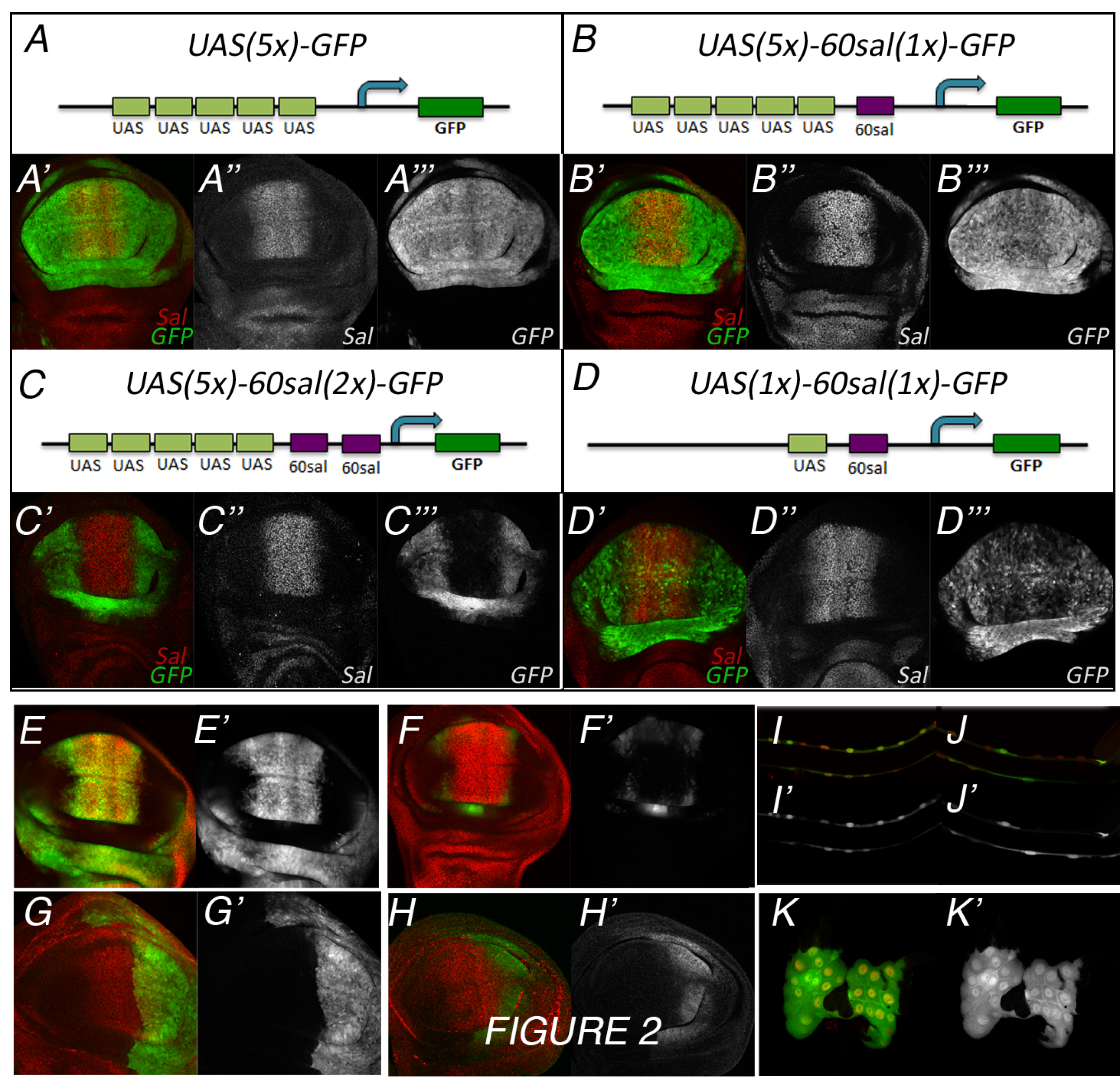
Characterization of the 60sal repressor element in heterologous constructs (A-A’’’) Schematic representation of the *UAS(5x)-GFP* construct (A) and expression of Salm (red in A’ and white in A’’) and GFP (green in A’ and white in A’’) in *nub-Gal4/ UAS(5x)-GFP* third instar wing discs. (B-B’’’) Schematic representation of the *UAS(5x)- 60sal(1x)-GFP* construct (B) and expression of Salm (red in B’ and white in B’’) and GFP (green in B’ and white in B’’) in *nub-Gal4/UAS(5x)-60sal(1x)-GFP* third instar wing discs. (C-C’’’) Schematic representation of the *UAS(5x)-60sal(2x)-GFP* construct (C) and expression of Salm (red in C’ and white in C’’) and GFP (green in C’ and white in C’’) in *nub-Gal4/UAS(5x)-60sal(2x)-GFP* third instar wing discs. (D-D’’’) Schematic representation of the *UAS(1x)-60sal(1x)-GFP* construct (D) and expression of Salm (red in D’ and white in D’’) and GFP (green in D’ and white in D’’) in *nub-Gal4/UAS(5x)- 60sal(2x)-GFP* third instar wing discs. (E-F) Expression of Salm (red) and GFP (green in E- F and white in E’-F’) in *sal-Gal4*/*UAS(5x)-GFP* (E-E’) and *sal-Gal4/UAS(5x)-60sal(2x)-GFP* (F-F’). (G-H) Expression of Salm (red) and GFP (green in G-H and white in G’-H’) in *hh- Gal4*/*UAS(5x)-GFP* (G-G’) and *sal-Gal4/UAS(5x)-60sal(2x)-GFP* (H-H’). (I-J) Expression of Salm (red) and GFP (green in I-J and white in I’-J’) in the trachea of *sal-Gal4*/*UAS(5x)-GFP* (I-I’) and *sal-Gal4/UAS(5x)-60sal(2x)-GFP* (J-J’). (K-K’) Expression of Salm (red) and GFP (green in I-J and white in I’-J’) in the ring gland of *sal-Gal4/UAS(5x)-60sal(2x)-GFP* (J-J’) larvae.

### The *60sal* repressive element is conserved in *Drosophilids*, and binds to Salm *in vitro* and *in vivo*

We next analyzed the possibility of direct binding of *in vitro* purified Salm with the *60sal* sequence. In these experiments (EMSA) we labeled the *60sal* sequence (^32^P-5’) and observed its migration in non-denaturing gels in the presence of different amounts of Salm (Fig. 3A). We observed a significant mobility shift starting at 1μg of Salm in the presence of Zn^2+^ salts (Fig. 3A). The presence of additional salts (NaCl, KCl and MgCl2) is, in general unfavorable for the interaction between Salm and *60sal* (Fig. 3A). The *60sal* sequence is very much conserved in different *Drosophilids*, and this conservation is particularly strong in the first 29 bp (Fig. 3B). We carried out EMSA assays with the 5’ and 3’ halves of the *60sal* sequence, and found that Salm binds to both fragments with similar efficiency (Fig 3C). Using the same experimental conditions (0.16 ng of labelled DNA and 150 nM ZnSO4) we found stronger retention for the *60sal* fragments than for its 5’ and 3’ halves, particularly when the amount of Salm is lower (0.75 mg). As a negative control, we assayed the possibility of binding between Salm and an independent sequence (*Sp1*), and found that Salm cannot bind to this sequence (Fig. 3C). To further analyze the contribution of the *60sal* to reporter expression *in vivo*, we generated new constructs containing three overlapping fragments of the *60sal* DNA (Δ5, Δ3.1 and Δ3.2; Fig. 3B) fused with 5 *UAS* sequences and the GFP coding sequence. In combinations with the *nub-Gal4* driver grown at 17°C we can detect a repressive effect of the *60sal* (Fig. 3D) and the 60salΔ5 (Fig. 3E) fragments, and much weaker repression in the cases of the 5’ 60salΔ3-1 (Fig. 3F) and 60salΔ3-2 fragments (Fig. 3G).

**Figure 3.**
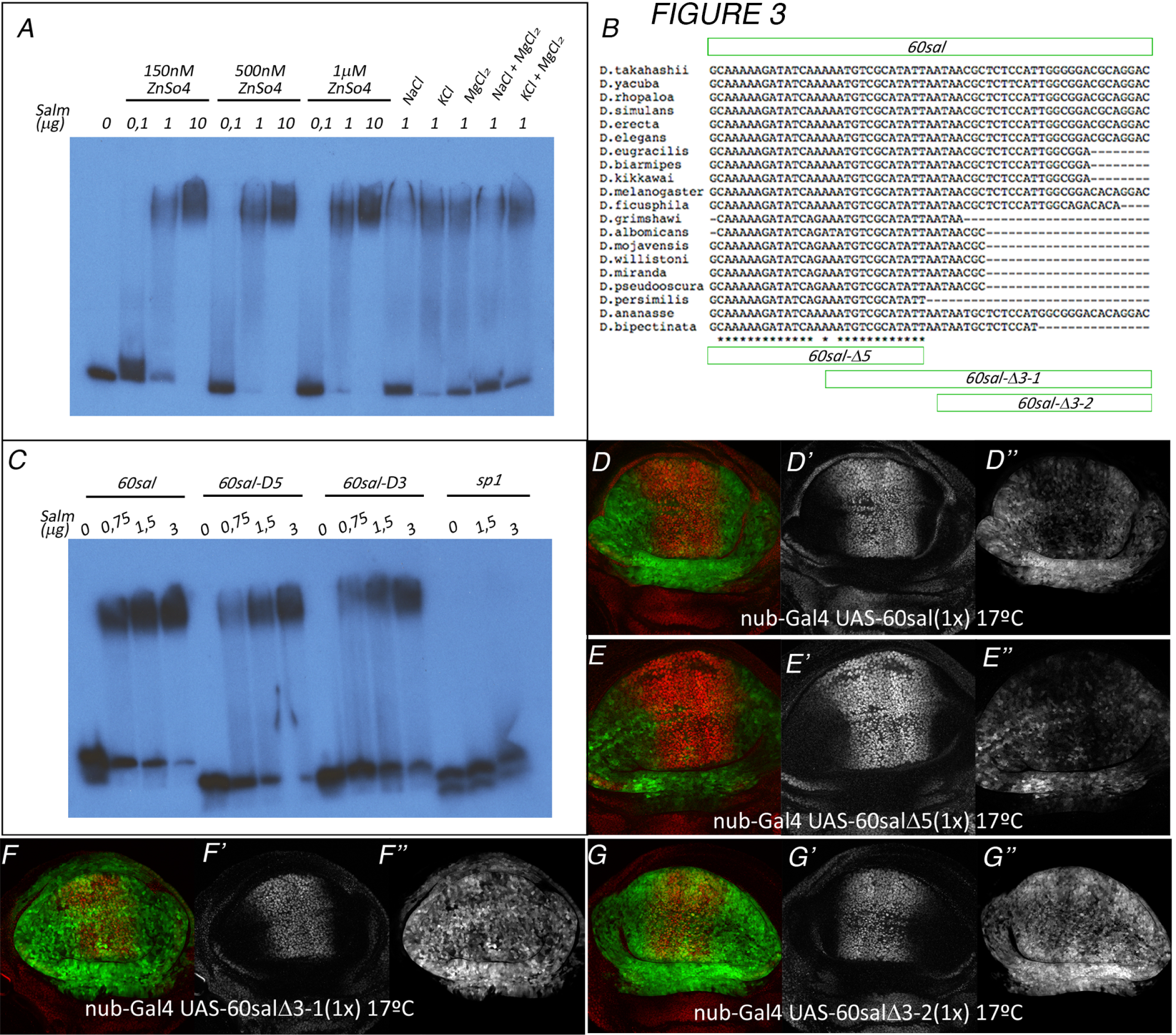
Binding of Salm to *60sal* in vitro (A) Electrophoretic mobility assays using Salm protein (0, 0.1μg, 1μg and 10μg) and 0,25μL (0,16ng) of the *60sal* double strand DNA fragment ^32^P-5’ labeled in the sense strain with γ^32^-ATP 50ng. (B) Sequence alignment of the *60sal* fragment in different *Drosophilids* (left column), and localization (green boxes) of the *60sal-*Δ*5*, *60sal-*Δ*3-1* and *60sal-*Δ*3.2* sub-fragments (below). (C) Electrophoretic mobility assays using Salm protein (0, 0.75μg, 1.5μg and 3μg) and 0,25μL (0,16ng) of *60sal*, *60sal-*Δ*5* and *60sal-*Δ*3-1* double strand DNA fragments whose sense strains are labelled with γ^32^-ATP 50ng. (D-D’’) Expression of Salm (red) and GFP in third instar wing discs of *nub-Gal4/UAS(5x)- 60sal(1x)-GFP* larvae grown at 17°C. The individual red and green channels are shown in D’ and D’’. (E-E’’) Expression of Salm (red) and GFP (green) in third instar wing discs of *nub-Gal4/UAS(5x)-60sal*Δ*5(1x)-GFP* larvae grown at 17°C. The individual red and green channels are shown in E’ and E’’. (F-F’’) Expression of Salm (red) and GFP (green) in third instar wing discs of *nub-Gal4/UAS(5x)-60sal*Δ*3-1(1x)-GFP* larvae grown at 17°C. The individual red and green channels are shown in F’ and F’’. (G-G’’) Expression of Salm (red) and GFP (green) in third instar wing discs of *nub-Gal4/UAS(5x)-60sal*Δ*3-2(1x)-GFP* larvae grown at 17°C. The individual red and green channels are shown in G’ and G’’.

Chromatin immunoprecipitation (ChIP) carried out with an endogenous Salm protein tagged with HA reveals a prominent peak of Salm binding in the same genomic position of the *60sal* sequence (Ostalé and de Celis, in preparation; Fig. 4A). This region, which includes the *sal60*Δ*5* fragment that retains most of the *60sal* repressor activity, is particularly enriched in A/T pairs (70%), the most prominent feature identified so far as Sal binding sequences (Barrio et al., 1996; Kong et al., 2021). We selected three 6 bp motives (AAATGT, CGCATA and TTAATA) located in the region of overlap between *60sal*Δ*5* and *60sal*Δ*3.1* and substituted all nucleotides independently for each motive to generate the reporter constructs *60sal-MUT1*, *60sal-MUT2* and *60sal-MUT3* (Fig. 4B). In all cases we observed that the repressor activity conferred by the *60sal* fragment (*5xUAS/2X60sal*) in combinations with the *nub-Gal4* driver is mostly lost (Fig. 4C-F). In this manner, the integrity of the second halve of the *60sal*Δ*5* sequence is absolutely required to confer Salm mediated repression, although this sequence, which is present in *60sal*Δ*3-1* is not sufficient for repression without the *60sal*Δ*5* first 18 nucleotides.

**Figure 4.**
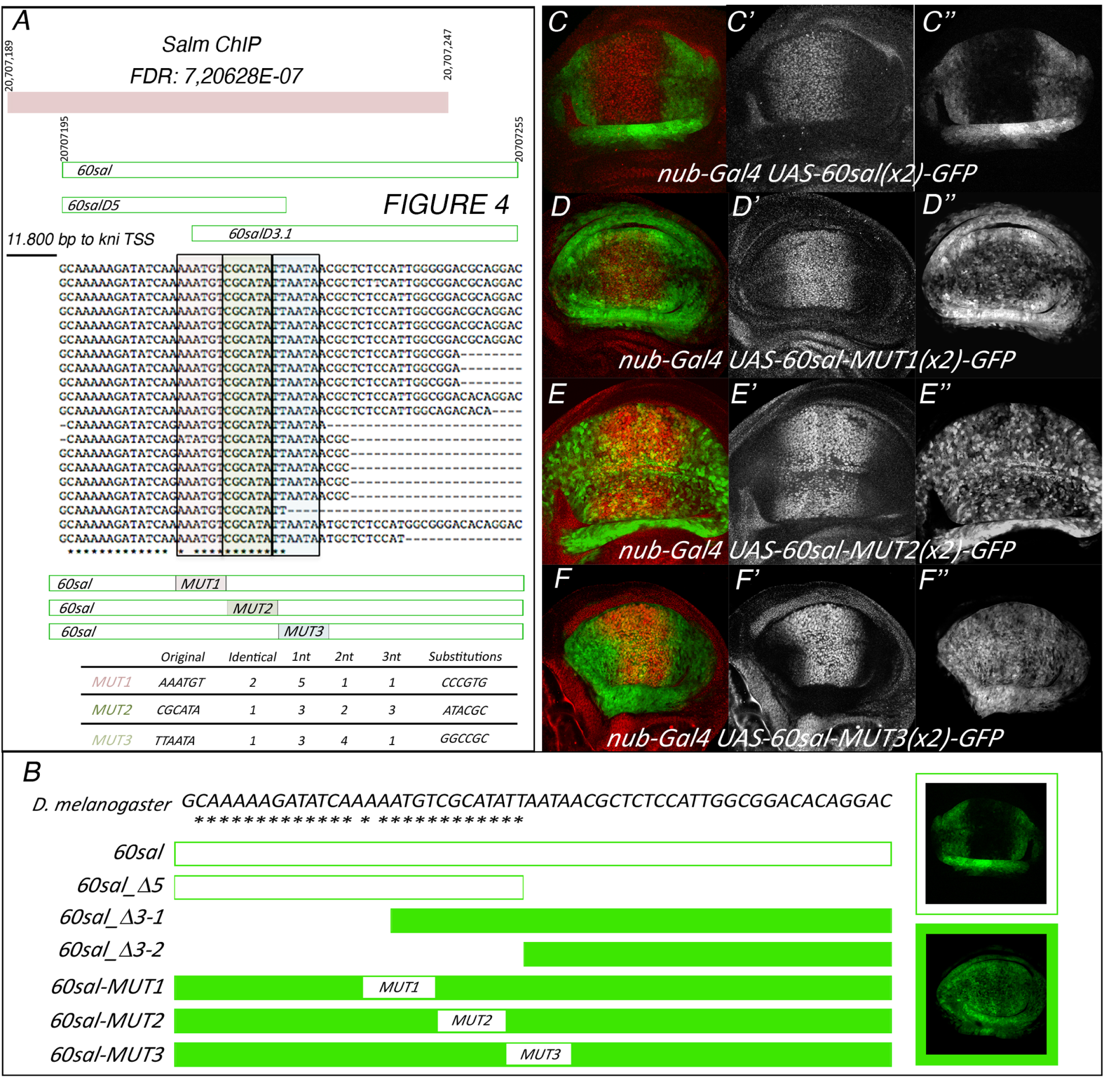
Chromatin immunoprecipitation of Salm identifies the 60sal repressor element. (A) Above: Schematic representation of the sequence isolated in Salm ChIP located at the *kni* locus. The genomic coordinates are indicated by numbers at the proximal and distal ends of the Salm ChIP and *60sal* fragments. Middle: Sequence alignment of the *60sal* fragment in different *Drosophilids* indicating the position and changes introduced in the *60sal* fragment (MUT1 to MUT3). (B) Sequence of the 60sal fragment and summary of the reporter expression results, where empty boxes indicate loss of reporter expression and filled boxes positive reporter expression in the central domain of the wing disc. (C-F) Expression of Salm (red) and GFP (green) in third instar wing discs of *nub- Gal4/UAS(5x)-60sal(2x)-GFP* (C-C’’), *nub-Gal4/UAS(5x)-60salMUT1(2x)-GFP* (D), *nub- Gal4/UAS(5x)-60salMUT2(2x)-GFP* (E) and *nub-Gal4/UAS(5x)-60salMUT3(2x)-GFP* larvae. The individual red and green channels are shown in C’-F’ and C’’-F’’, respectively.

Our analysis identifies a minimal 30bp sequence, *60sal-*Δ*5*, conserved in all *Drosophilids* that displays Salm binding *in vivo* and *in vitro* and that mediates Salm/Salr transcriptional repression in the central region of the wing blade. To our knowledge, this is the first case in which a regulatory interaction between the Salm protein and one of its targets in *Drosophila* is shown to rely on Salm binding to a target sequence, and confirm that in this instance, the Salm protein acts as a canonical sequence-specific DNA binding protein with transcriptional repressor activity.

## Acknowledgments

We thank Rosa Barrio, Carlos Estella, Gerardo Jiménez and Frank Schnorrer for reagents and Ana Ruiz-Gómez for help in the purification of Salm protein. We also thank the Developmental Studies Hybridoma Bank at Iowa University, NIG-FLy in Japan, Bloomington Stock Center and VDRC Stock Center for providing the tools necessary for this work. We would also like to acknowledge the support from the *Drosophila* transgenesis and confocal microscopy CBMSO scientific services. The CBMSO enjoys institutional support from the Ramón Areces Foundation. This research was supported by Secretaría de Estado de Investigación, Desarrollo e Innovación, Grant/Award Number PGC2018-094476-B-I00. The founders have no role in the research design, execution, analysis or interpretation of the results. The authors affirm that all data necessary for confirming the conclusions of the article are present within the article, figures, tables and supplementary information.

---

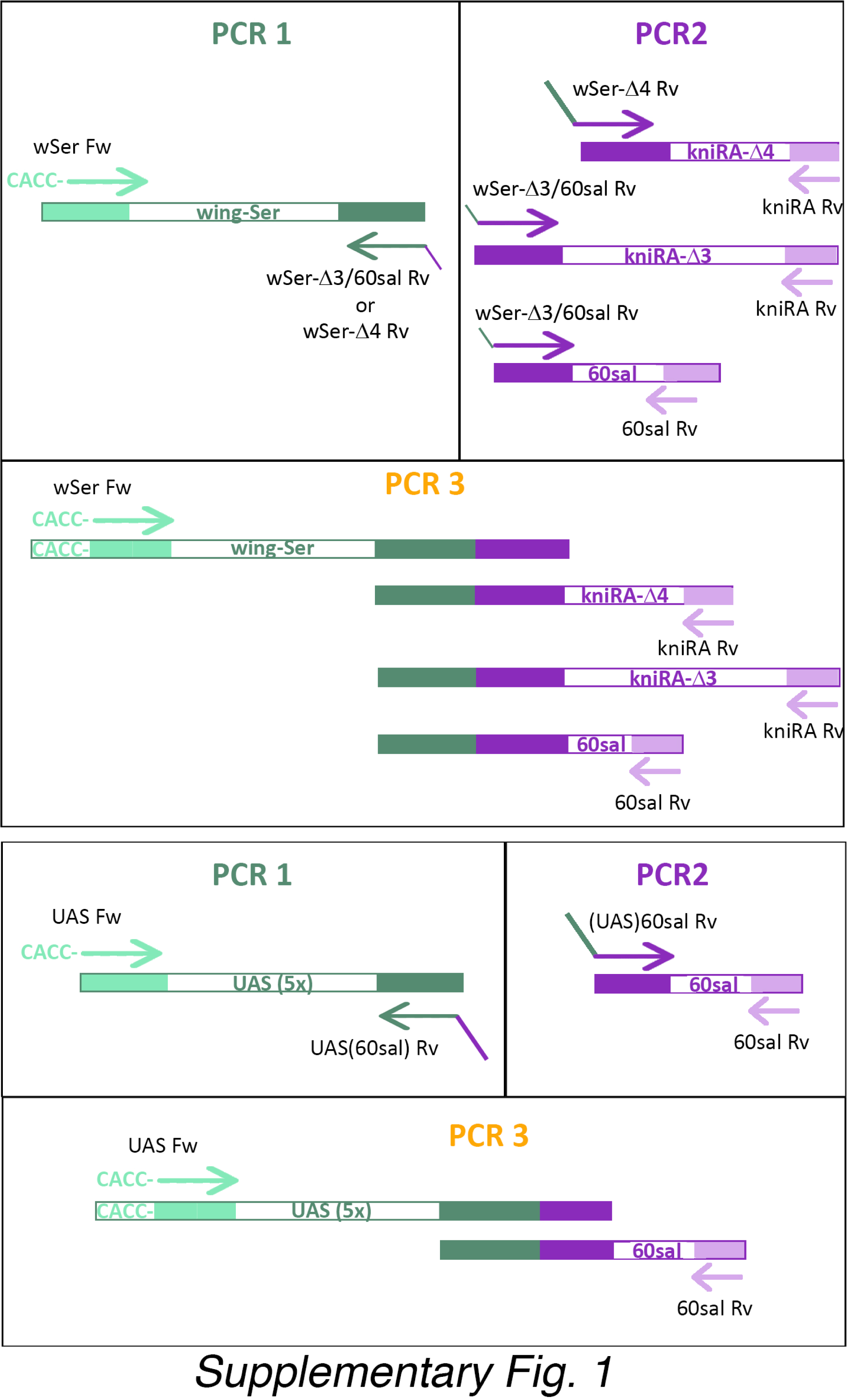

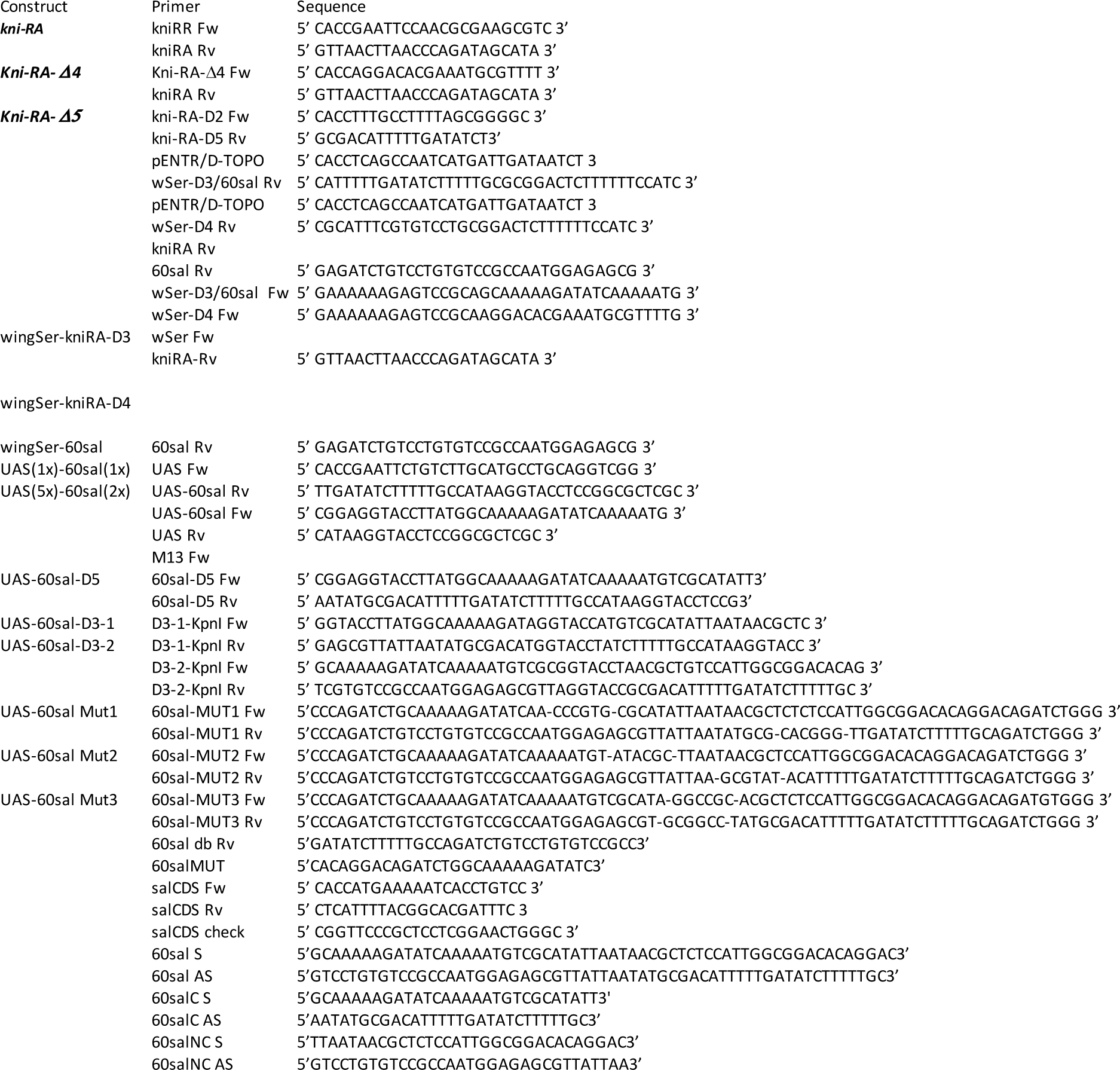

## LITERATURE CITED

1. Alvarez, C., Quiroz, A., Benitez-Riquelme, D., Pincheira, R., Riffo, E. and Castro, A.F. (2021). SALL proteins; common and antagonistic roles in cancer. Cancers 13, 6292. DOI: 10.3390/CANCERS13246292.

2. Barrio R, Shea MJ, Carulli J, Lipkow K, Gaul U, Frommer G, Schuh R, Jäckle H, Kafatos FC. (1996). The *spalt-related* gene of *Drosophila melanogaster* is a member of an ancient gene family, defined by the adjacent, region-specific homeotic gene *spalt*. Dev Genes Evol. 1996 Dec;206(5):315–25. doi: 10.1007/s004270050058.

3. Barrio R,de Celis JF, Bolshakov S and Kafatos FC (1999). Identification of regulatory regions driving the expression of the Drosophila spalt complex at different developmental stages. Dev. Biol. 215: 33–47. DOI: 10.1006/dbio.1999.9434.

4. Bischof J, Maeda RK, Hediger M, Karch F, Basler K. (2007). An optimized transgenesis system for Drosophila using germ-line-specific phiC31 integrases. Proc Natl Acad Sci USA 104(9):3312–3317. doi: 10.1073/pnas.0611511104.

5. Borozdin W, Boehm D, Leipoldt M, Wilhelm C, Reardon W, Clayton-Smith J, Becker K, Mühlendyck H, Winter R, Giray O, Silan F and Kohlhase J (2004). SALL4 deletions are a common cause of Okihiro and acro-renal-ocular syndromes and confirm haploinsufficiency as the pathogenic mechanism. J Med Genet 41:e113. DOI: 10.1136/jmg.2004.019901.

6. Boy AL, Zhai Z, Habring-Müller A, Kussler-Schneider Y, Kaspar P and Lohmann I. (2010). Vectors for efficient and high-throughput construction of fluorescent *Drosophila* reporters using the PhiC31 site-specific integration system. Genesis 48: 452–456. DOI: 10.1002/dvg.20637.

7. Calleja M, Moreno E, Pelaz S, Morata G. (1996). Visualization of gene expression in living adult *Drosophila*. Science. 274:252–255. doi: 10.1126/science.274.5285.252.

8. Cantera R, Lüer K, Rusten TE, Barrio R, Kafatos FC, Technau GM. (2002). Mutations in *spalt* cause a severe but reversible neurodegenerative phenotype in the embryonic central nervous system of *Drosophila melanogaster*. Development 129:5577–5586. doi: 10.1242/dev.00158.

9. Carthew RW, Chodosh LA, Sharp PA (1985) An RNA polymerase II transcription factor binds to an upstream element in the adenovirus major late promoter. Cell 43: 439–448. DOI: 10.1016/0092-8674(85)90174-6.

10. Cruz C, Glavic A, Casado M, de Celis JF. (2009). A gain-of-function screen identifying genes required for growth and pattern formation of the Drosophila melanogaster wing. Genetics 183(3):1005–1026. doi: 10.1534/genetics.109.107748.

11. de Celis, J., Barrio, R. and Kafatos, F. (1996). A gene complex acting downstream of *dpp* in *Drosophila* wing morphogenesis. Nature 381, 421–424. DOI: 10.1038/381421a0.

12. de Celis, J.F., Barrio, R., Kafatos, F.C. (1999). Regulation of the *spalt/spalt-related* gene complex and its function during sensory organ development in the *Drosophila* thorax. Development 126(12): 2653–2662. DOI: 10.1242/dev.126.12.2653.

13. de Celis JF, Barrio R. (2000). Function of the *spalt/spalt-related* gene complex in positioning the veins in the *Drosophila* wing. Mech Dev. 91:31–41. doi: 10.1016/s0925-4773(99)00261-0.

14. de Celis J F and Barrio R (2009). Regulation and function of Spalt proteins during animal development. Int. J. Dev. Biol. 53: 1385–1398. DOI: 10.1387/ijdb.072408jd.

15. Grieder, NC, Morata G, Affolter M and Gehring WJ (2009). Spalt major controls the development of the notum and of wing hinge primordia of the *Drosophila melanogaster* wing imaginal disc. Dev. Biol. 329: 315–326. DOI: 10.1016/j.ydbio.2009.03.006

16. Hu N and Castelli-Gair, J (1999). Study of the posterior spiracles of *Drosophila* as a model to understand the genetic and cellular mechanisms controlling morphogenesis. Dev. Biol. 214: 197–210. DOI: 10.1006/dbio.1999.9391.

17. Kohlhase, J (2000). SALL1 mutations in Townes-Brocks syndrome and related disorders. HUMAN MUTATION 16: 460–466. DOI: 10.1002/1098-1004.

18. Kong NR, Bassal MA, Tan HK, Kurland JV, Yong KJ, Young JJ, Yang Y, Li F, Lee JD, Liu Y, et al. (2021). Zinc finger protein SALL4 functions through an AT-rich motif to regulate gene expression. Cell Rep 34: 108574. 10.1016/j.celrep.2020.108574.

19. Kosman D, Small S and Reinitz J. (1998). Rapid preparation of a panel of polyclonal antibodies to Drosophila segmentation proteins. Dev. Genes Evol. 208, 290–294. DOI: 10.1007/s004270050184.

20. Kühnlein RP and Schuh R. (1996). Dual function of the region-specific homeotic gene *spalt* during *Drosophila* tracheal system development. Development 122(7):2215–2223. doi: 10.1242/dev.122.7.2215.

21. Lunde K, Biehs B, Nauber U and Bier E. (1998). The *knirps* and *knirps-related* genes organize development of the second wing vein in *Drosophila*. Development 125: 4145–4154. DOI: 10.1242/dev.125.21.4145.

22. Lunde K, Trimble JL, Guichard A, Guss KA, Nauber U, Bier E. (2003). Activation of the knirps locus links patterning to morphogenesis of the second wing vein in Drosophila Development 130: 235–248. 10.1242/dev.00207.

23. Martin M, Organista MF and de Celis JF (2014). Structure of developmental gene regulatory networks from the perspective of cell fate-determining genes. Transcription 7: 32–37. DOI: 10.1080/21541264.2015.1130118.

24. Martin M, Ostalé CM, de Celis JF (2017). Patterning of the *Drosophila* L2 vein is driven by regulatory interactions between region-specific transcription factors expressed in response to Dpp signalling. Development 144: 3168–3176. 10.1242/dev.143461.

25. Nikki R. Kong, Mahmoud A. Bassal, Hong Kee Tan, Jesse V. Kurland, Kol Jia Yong, John J. Young, Yang Yang, Fudong Li, Jonathan D. Lee, Yue Liu, Chan-Shuo Wu, Alicia Stein, Hongbo R. Luo, Leslie E. Silberstein, Martha L. Bulyk, Daniel G. Tenen, Li Chai (2021). Zinc Finger Protein SALL4 Functions through an AT-Rich Motif to Regulate Gene Expression. Cell Reports 34 108574. DOI: 10.1016/j.celrep.2020.108574.

26. Organista MF and de Celis JF (2012) The Spalt transcription factors regulate cell proliferation, survival and epithelial integrity downstream of the Decapentaplegic signalling pathway. Biology Open 2, 37–48. doi: 10.1242/bio.20123038.

27. Powell CM, Michaelis RC (1999). Townes-Brocks syndrome. Journal of Medical Genetics 36: 89–93. DOI: 10.1136/jmg.36.2.89.

28. Ru W, Koga T, Wang X, Guo Q, Gearhart MD, Zhao S, Murphy M, Kawakami H, Corcoran D, Zhang J, Zhu Z, Yao X, Kawakami Y, Xu C. (2022). Structural studies of SALL family protein zinc finger cluster domains in complex with DNA reveal preferential binding to an AATA tetranucleotide motif. J Biol Chem. 298(12):102607. doi: 10.1016/j.jbc.2022.102607.

29. Rusten TE, Cantera R, Urban J, Technau G, Kafatos FC, Barrio R. (2001). Spalt modifies EGFR-mediated induction of chordotonal precursors in the embryonic PNS of Drosophila promoting the development of oenocytes. Development 128:711–722. doi: 10.1242/dev.128.5.711.

30. Schönbauer C, Distler J, Jährling N, Radolf M, Dodt HU, Frasch M, Schnorrer F. (2011). Spalt mediates an evolutionarily conserved switch to fibrillar muscle fate in insects. Nature 479(7373):406–409. doi: 10.1038/nature10559.

31. Shea MJ, King DL, Conboy MJ, Mariani BD, Kafatos FC (1990). Proteins that bind to Drosophila chorion cis-regulatory elements: a new C2 H2 zinc finger protein and a C2C2 steroid receptor-like component. Genes Dev 4: 1128–1140.

32. Sweetman D and Münsterberg A. (2006). The vertebrate *spalt* genes in development and disease. Dev. Biol. 293: 285–293. DOI: 10.1016/J.YDBIO.2006.02.009.

33. Yan SJ, Gu Y, Li WX and Fleming RJ. (2004). Multiple signaling pathways and a selector protein sequentially regulate Drosophila wing development. Development 131(2): 285–298. DOI: 10.1242/dev.00934.

34. Xu X-R, Gantz VM, Siomava N and Bier E. (2017). CRISPR/Cas9 and active genetics-based trans-species replacement of the endogenous Drosophila kni-L2 CRM reveals unexpected complexity. eLife 6: e30281. DOI: 10.7554/eLife.30281.

